# Analysis of clinical *Bordetella pertussis* isolates using whole genome sequences reveals novel genomic regions associated with recent outbreaks in the United States of America

**DOI:** 10.1101/047886

**Authors:** Glen Otero, Benjamin M. Althouse, Samuel V. Scarpino

**Author notes:** Benjamin M. Althouse, PhD, ScM Institute for Disease Modeling 3150 139th Ave SE, Bellevue, WA, 98118 Santa Fe Institute, Santa Fe, NM Department of Biology, New Mexico State University, Las Cruces, NM. Samuel V. Scarpino, PhD Department of Mathematics and Statistics, University of Vermont 16 Colchester Ave., Burlington, VT 05401 Santa Fe Institute, Santa Fe, NM Complex Systems Center, University of Vermont, Burlington, VT. These authors contributed equally to this work.

## Abstract

**Background:** Despite high-levels of vaccination, whooping cough, primarily caused by Bordetella pertussis (BP), has persisted and resurged. It remains a major cause of infant death worldwide and is the most prevalent vaccine-preventable disease in developed countries. To date, most genomic studies have focused on a small subset of the BP genome, biasing our clinical understanding and public health awareness.

**Methods:** We performed a Genome-Wide Association Study (GWAS) on 76 U.S. BP whole genomes, including strains from recent outbreaks.

**Results:** A GWAS of the 76 BP isolates revealed a sharp increase in genetic variation associated with the Minnesota 2012 outbreak and identified 52 variants unique to the Minnesota outbreak and 19 unique to the California and Washington outbreaks. None of the identified variants were shared between the outbreaks and the vast majority were previously uncharacterized. We further identified variation associated with pertactin negative strains and acellular vaccination.

**Conclusions:** We identified novel genomic regions associated with recent BP outbreaks. Our results underscore the need for increased whole genome sequencing of BP isolates, which can reduce costly misdiagnosis and improve surveillance. The genes containing these variants warrant further investigation into their possible roles in BP pathogenicity and the ongoing resurgence in the U.S.

## Introduction

The ongoing whooping cough epidemic - caused primarily by *Bordetella pertussis* - is rapidly becoming a public health emergency in the United States of America (U.S.) [1]. The recent outbreaks in 2012 generated the highest number of whooping cough cases observed since the pre-vaccine era [2], and, in 2014, California had the most cases ever recorded in the state [3].

Clinicians rely on genetic-based tests to rapidly and accurately determine whether an individual is infected with *B. pertussis* and public health officials use genetic results to monitor outbreaks and the effectiveness of interventions [4, 5]. However, despite advances in next generation sequencing (NGS) technology and expanded specimen collection, the majority of *B. pertussis* genetic studies focus on detecting various genetic markers associated with only a handful of known virulence factors and vaccine components [6, 7]. This approach is unable to identify other genomic regions involved in *B. pertussis* pathogenicity and/or vaccine resistance potentially responsible for clonal expansion and persistence in a vaccinated population.

To investigate the clinical and public health consequences of using fixed regions of the *B. pertussis* genome for diagnosis and reporting, we conducted a genome wide association study (GWAS) on *B. pertussis* genomes isolated during three recent U.S. outbreaks. We detected an increase in genetic variation in the Minnesota 2012 subpopulation and 31 non-synonymous single nucleotide polymorphisms (SNPs) not associated with other outbreaks. These findings indicate that it is possible to differentiate *B. pertussis* subpopulations underlying outbreaks and track the spread of the bacteria using genomic data. The same methods were used to identify SNPs associated with pertactin(-) strains and strains isolated during the acellular vaccine period (1998-2012). Importantly, the identified SNPs are not located in genes currently associated with known virulence or vaccine components and thus represent an opportunity for the design of novel therapeutics.

These findings broaden our understanding of the evolutionary forces acting on *B. pertussis*, the genetic loci undergoing selective pressure and how *B. pertussis* may be evolving in response to vaccine pressure.

## Methods

### Strains studied

We utilized genomic data from 76 *B. pertussis* strains isolated in the U.S. between 1935-2012. Thirty five strains were collected from various locations in the U.S. between 1935-2005 (referred to as the historical strains), 19 strains were collected from the 2010 California outbreak [8], 9 strains were collected from the 2012 Washington outbreak [9], 12 strains were collected from the 2012 Minnesota outbreak [10], and *B. pertussis* strain 18323, isolated in the U.S. in 1946 and for which a detailed Genbank file is available, was the reference genome used for comparison. Except for the Minnesota outbreak genome sequences, all genome sequence data was downloaded from the NCBI Sequence Read Archive. NCBI Sequence Read Archive accession numbers for the genome sequences are listed in eTable 1 in the Supplement. The Minnesota outbreak whole genome sequence data were obtained from the researchers who isolated and sequenced the Minnesota outbreak strains [10].

**Table 1:** Non-synonymous and Nonsense SNPs significantly associated with the Minnesota outbreak strains. Gene and protein annotations are from the 18323 reference genome Genbank file.

| Gene | Protein | SNP Type | Codon position | Reference amino acid | Non-reference amino acid |
| --- | --- | --- | --- | --- | --- |
| tsr | methyl-accepting chemotaxis protein I | Non-synonymous | 498 | Ala | Val |
| putA | bifunctional proline oxidoreductase/transcriptional repressor | Non-synonymous | 1222 | Ala | Val |
| nuoD | respiratory-chain NADH dehydrogenase, 49 kDa subunit | Non-synonymous | 313 | Ile | Ser |
| mutL | DNA mismatch repair protein | Non-synonymous | 356 | Ala | Val |
| maeB | NADP-dependent malic enzyme | Non-synonymous | 628 | Gly | Ser |
| maeB | NADP-dependent malic enzyme | Non-synonymous | 628 | Gly | Asp |
| maeB | NADP-dependent malic enzyme | Non-synonymous | 625 | Gln | Arg |
| maeB | NADP-dependent malic enzyme | Non-synonymous | 643 | Ser | Thr |
| lnt | putative apolipoprotein N-acyltransferase | Non-synonymous | 15 | Ile | Thr |
| hmp1 | ribonuclease E | Non-synonymous | 653 | Thr | Ala |
| fliK | flagellar hook-length control protein | Non-synonymous | 338 | Ala | Thr |
| dadA | D-amino acid dehydrogenase small subunit | Non-synonymous | 258 | Arg | His |
| ctaD | cytochrome c oxidase polypeptide I | Non-synonymous | 239 | Thr | Ala |
| carB | carbamoyl-phosphate synthase large chain | Non-synonymous | 796 | Ser | Gly |
| argD | putative acetylornithine aminotransferase | Non-synonymous | 111 | Gly | Ala |
| BN118_3679 | putative inner membrane protein | Non-synonymous | 51 | Leu | Pro |
| BN118_3610 | TolA protein / Proline-rich inner membrane protein | Non-synonymous | 237 | Ala | Pro |
| BN118_3610 | TolA protein / Proline-rich inner membrane protein | Non-synonymous | 237 | Ala | Glu |
| BN118_3090 | conserved hypothetical protein | Non-synonymous | 65 | Val | Leu |
| BN118_2919 | probable enoyl-CoA hydratase/isomerase | Non-synonymous | 6 | Ser | Leu |
| BN118_2793 | conserved hypothetical protein | Non-synonymous | 264 | Ala | Thr |
| BN118_2781 | conserved hypothetical protein | Non-synonymous | 176 | Leu | Phe |
| BN118_2428 | ABC transport protein, inner membrane component | Non-synonymous | 110 | Arg | His |
| BN118_2296 | putative membrane protein | Non-synonymous | 75 | Pro | Gln |
| BN118_2252 | putative outer membrane protein | Non-synonymous | 481 | Pro | Ser |
| BN118_1681 | probable LysR-family transcriptional regulator | Non-synonymous | 160 | Asp | Glu |
| BN118_0965 | putative permease component of branched-chain amino acid transport system | Non-synonymous | 454 | Gly | Asp |
| BN118_0794 | putative periplasmic protein | Non-synonymous | 199 | Gly | Asp |
| BN118_0462 | putative exported protein | Non-synonymous | 154 | Ala | Val |
| BN118_0138 | putative exported protein | Non-synonymous | 69 | Ala | Glu |
| BN118_3321 | conserved hypothetical protein | Nonsense | 163 | Gln | STO |

### Whole genome sequencing and phylogenetic analysis

Short read sequences from each of the *B. pertussis* genomes underwent quality checks with FastQC, had adapters trimmed with Trimmomatic, were mapped to the *B. pertussis* reference genome 18323 with bowtie2 and SNPs called using bcftools. Parsnp [11] was used to further filter the SNPs to produce 6079 core-genome SNPs, create a whole genome alignment and maximum likelihood phylogenetic tree of the 76 *B. pertussis* genomes. Given the very high support for the tree and all internal nodes, we do not discuss the results of a Bayesian phylogenetic analysis as the posterior distribution of trees was tightly peaked around the maximum likelihood tree.

### Genome wide association study

In order to identify genomic regions significantly associated with the various outbreaks we conducted a GWAS on the 6079 core-genome SNPs following established best practices [12]. The 6079 SNPs from each genome were concatenated and an association between each SNP and the phenotype was tested by logistic regression implemented in R [12]. Each of the *B. pertussis* groups, e.g. Minnesota, California, Washington, and historical, were treated as a phenotype and the rest of the genomes used as controls. Due to their similar number of SNPs/genome and phylogenetic similarity we combined the California and Washington outbreak strains into a single phenotype while using the remaining genomes as controls. Strain 18323 was used as the reference genome and its Genbank file used to annotate the SNPs. Multiple testing was accounted for by applying a Bonferroni correction. The individual locus effect of a SNP was considered significant if its p-value was smaller than *α*/*n*, with *α* = 0.05 as the genome-wide false positive rate and n is the number of SNPs. The genome-wide –log_10_P value value threshold was 5.0.

## Results

### *B. pertussis* genome SNP frequencies in the US from 1935-2012

To investigate possible SNP accumulation trends of *B. pertussis* genomes isolated in the U.S., we calculated the median number of SNPs/genome for each year we had *B. pertussis* whole genome sequence data. Figure 1 illustrates that the number of SNPs/genome fluctuated very little each year around a median of 1940 SNPs/genome until 2012 when there was a 22% increase to 2367 SNPs/genome. Due to the large range of SNPs/genome values in 2012, we separated the strains from 2012 into two groups, those isolated in the Washington outbreak and those isolated in the Minnesota outbreak. We also grouped the remaining genomes isolated between 1935-2005 by decade.

**Figure 1:**
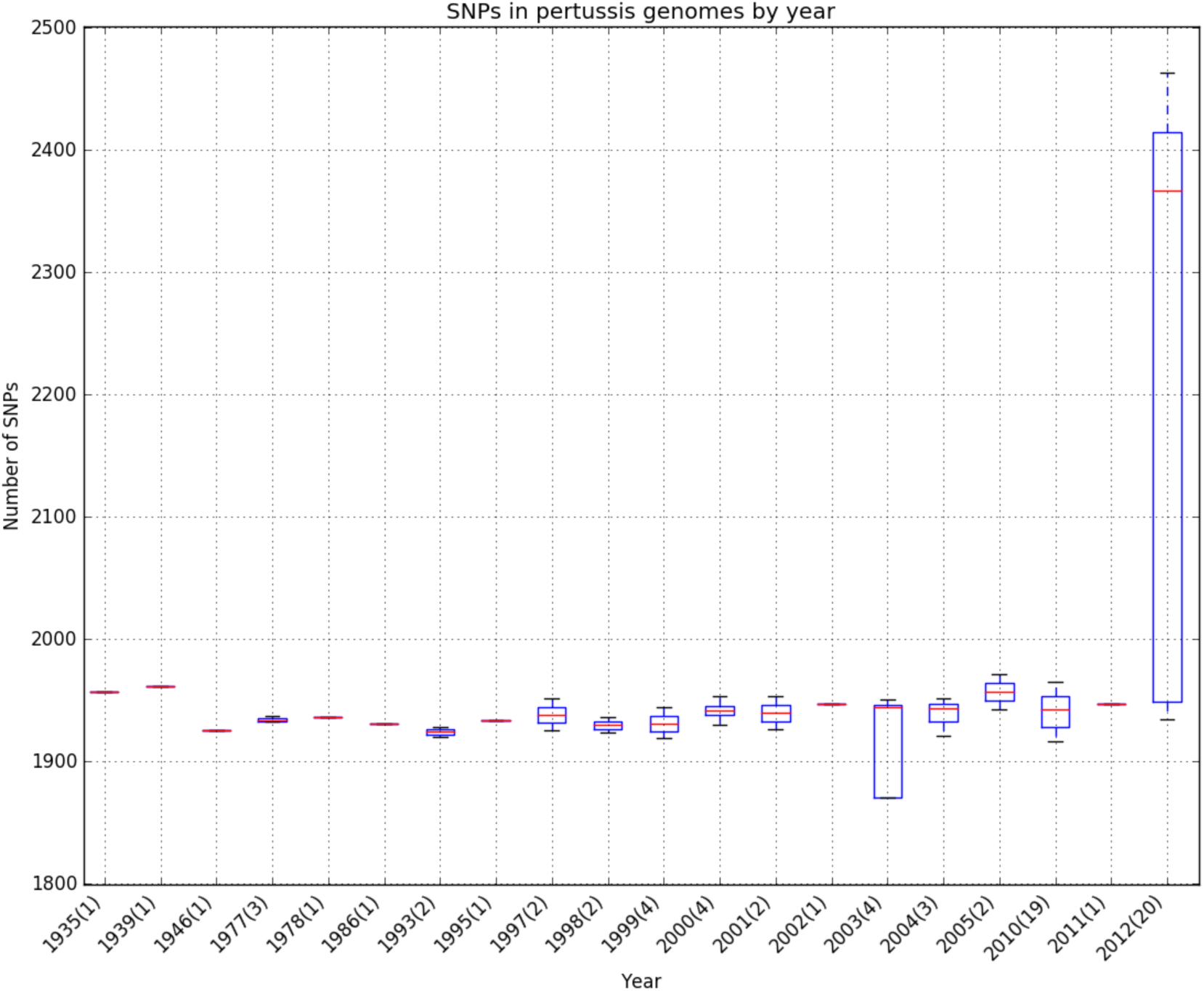
Median SNPs/genome in all *B. pertussis* strains. Boxplots show median SNPs/genome studied grouped by year. Numbers in parentheses indicate number of genomes for the given year.

Separating the 2012 strains geographically revealed that the Minnesota strains were the source of the increase in 2012 SNP counts with a median 2408 SNPs/genome, 24% more SNPs/genome than the 1943 SNPs/genome median of all other periods, including those from the California 2010 (1942 SNPs/genome) and Washington 2011 & 2012 (1947 SNPs/genome) isolates (Figure 2). It is unlikely the increased SNPs/genome in the Minnesota strains can be attributed to imbalances in genome coverage that favor finding more SNPs in the Minnesota genomes as the depth of coverage is 49x for the Minnesota genomes, 90x for the California and Washington genomes and 60x for the historical genomes. Likewise, the breadth of coverage is 90% for the Minnesota genomes, 88% for the California and Washington genomes and 88% for the historical genomes.

**Figure 2:**
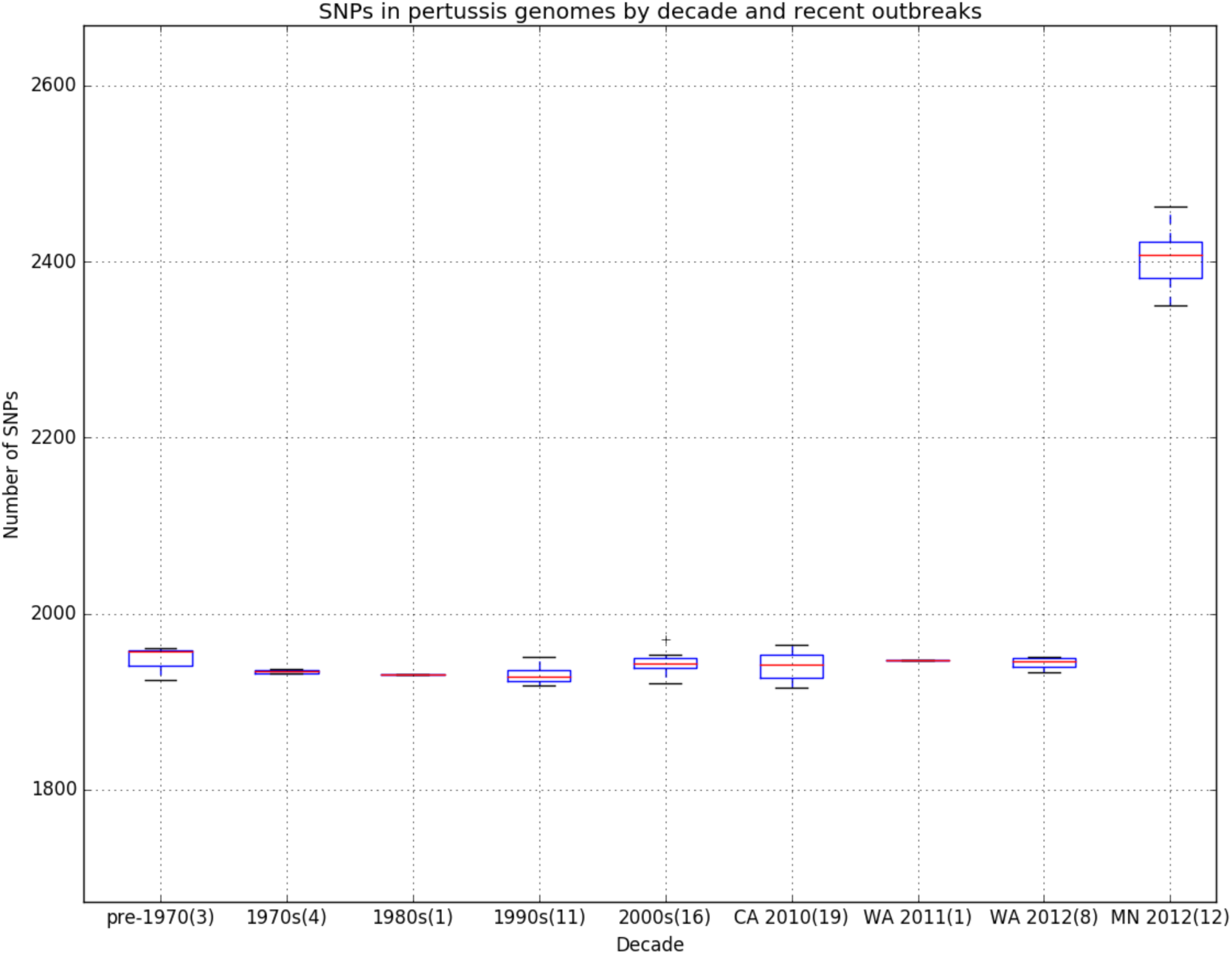
Median SNPs/genome in all *B. pertussis* strains. Boxplots show median SNPs/genome studied grouped by decade and by outbreak. Numbers in parentheses indicate number of genomes for the given decade/outbreak.

### Phylogenetic analysis of *B. pertussis* strains

Greater than 99% of each genome was included in the whole genome alignment that was used to generate the phylogenetic tree in Figure 3. The genomes cluster roughly in chronological order with a few exceptions. Twenty four of the 28 California and Washington strains cluster together. Three of the four California strains are in a clade that contains the 18323 reference genome that was isolated in 1946. One of the three California strains that clusters with 18323 differs by only seven SNPs, the other two each by 22 SNPs. This cluster of three California strains with 18323 is evidence that a subpopulation of *B. pertussis* very similar to strains present 70 years ago are still in circulation and causing disease.

**Figure 3:**
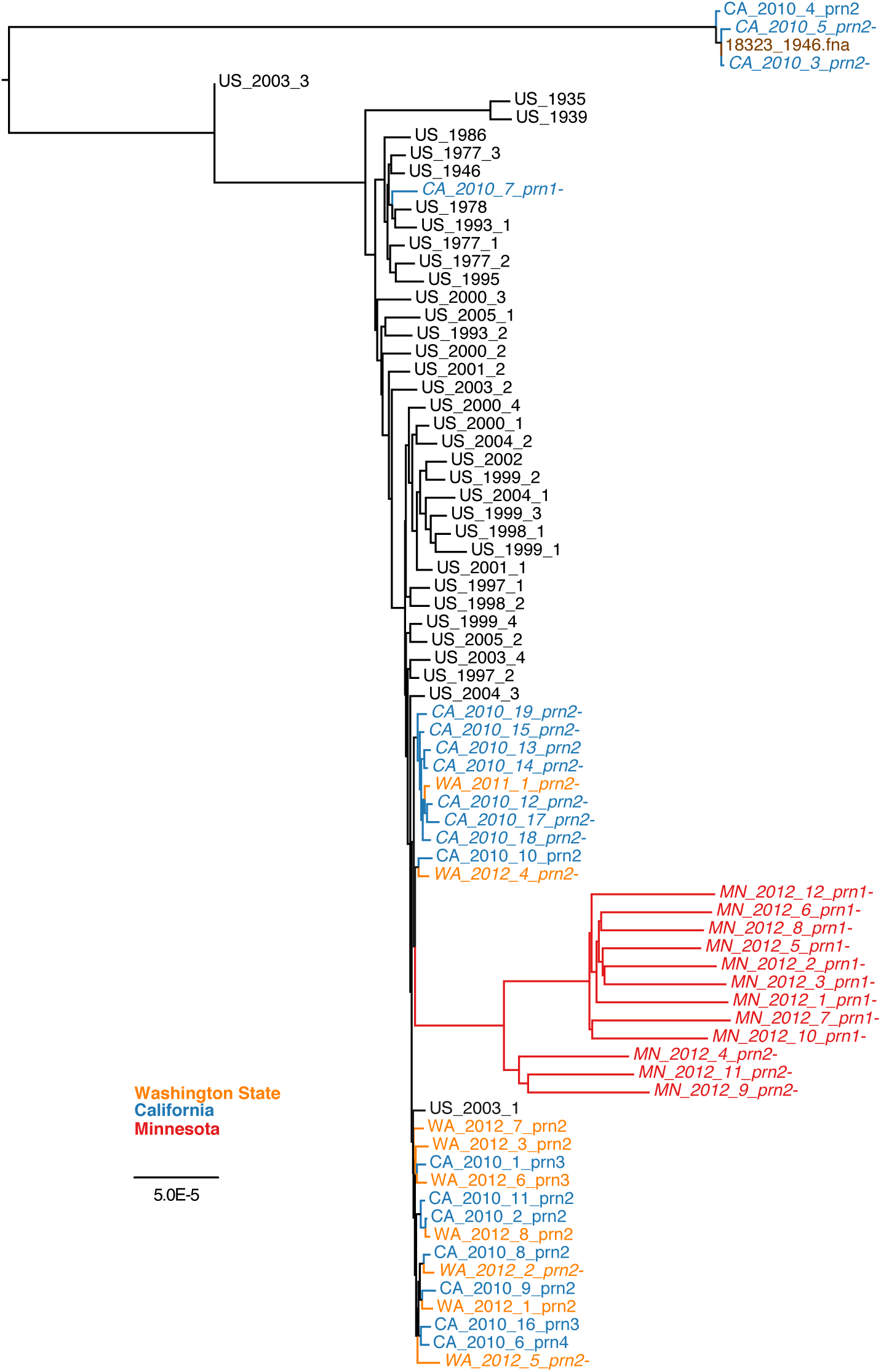
Phylogenetic tree of 76 *B. pertussis* genomes based on whole genome alignment. Genomes are color-coded: black indicates historical genomes; red, Minnesota outbreak; orange, Washington State outbreak; blue, California outbreak; and brown, reference genome. The Minnesota, California, and Washington outbreak genomes are labeled with their state, year, sample number, and pertactin allele. If the pertactin gene was disrupted, there is a - sign after the pertactin allele and the label is italicized. The unit of scale is substitutions/site.

The 12 strains isolated in the Minnesota outbreak form a distinct clade with two distinct subclusters. One subcluster contains nine genomes that lack a pertactin (prn1) signal sequence and the other cluster contains three genomes in which pertactin (prn2) is disrupted by insertion sequence IS481 or contains a stop codon. The majority of the pertactin(-) strains studied cluster around the Minnesota clade except for three pertactin(-) strains from the California outbreak found in the upper half of the tree. This suggests that pertactin(-) strains have arisen independently multiple times in the U.S., evidence for which has been previously reported [13]. Two of these pertactin(-) strains cluster with 18323, indicating that *B. pertussis* strains very similar to those present 70 years ago are still in circulation and have become pertactin(-). The phylogenetic tree in Figure 3 illustrates the uniqueness of the Minnesota subpopulation and the 18323-associated cluster of CA strains compared to the rest of the genomes studied.

### Polymorphisms characteristic of three recent U.S. outbreaks of *B. pertussis*

Out of 6079 SNPs from the 76 *B. pertussis* strains we identified 52 SNPs in coding regions that were significantly associated (–log_10_P value ≥ 5) with the Minnesota outbreak. Of the 52 SNPs, the 31 non-synonymous or nonsense SNPs characteristic of the Minnesota subpopulation are listed in Table 1.

The Washington outbreak GWAS did not identify any significantly associated SNPs. However, due to their similar number of SNPs/genome and phylogenetic similarity, we combined the California and Washington outbreak strains into a single phenotype while using the remaining genomes as controls in a GWAS. From 6079 SNPs we identified 19 SNPs in coding regions that were significantly associated (–log_10_P value ≥ 5) with the combined California/Washington outbreak strains. The 19 SNPs are listed in Table 2. The California/Washington SNPs overlapped with nine SNPs of the historical strains suggesting that some of the California/Washington strains are more closely related to the historical strains. This is supported by the way these strains cluster in the phylogenetic tree.

**Table 2:** SNPs significantly associated with the California and Washington outbreaks. Gene and protein annotations are from the 18323 reference genome Genbank file.

| Gene | Protein | SNP Type | Codon position | Reference amino acid | Non-reference amino acid |
| --- | --- | --- | --- | --- | --- |
| tsr | methyl-accepting chemotaxis protein I | Synonymous | 184 | Ala | Ala |
| bfrI | putative ferrisiderophore receptor | Non-synonymous | 219 | Arg | Gln |
| BN118_2634 | threonine synthase | Non-synonymous | 337 | Glu | Gly |
| BN118_2561 | hypothetical protein | Non-synonymous | 135 | Val | Gly |
| BN118_2262 | putative phospholipase | Non-synonymous | 120 | Ala | Gly |
| BN118_2256 | putative transferase | Non-synonymous | 362 | Ala | Gly |
| BN118_2154 | putative ABC-transport protein, ATP-binding component | Non-synonymous | 70 | His | Pro |
| BN118_1839 | putative oxidoreductase | Synonymous | 43 | Arg | Arg |
| BN118_1806 | probable probable aldehyde dehydrogenase | Non-synonymous | 88 | Ala | Gly |
| BN118_0965 | putative permease component of<br>branched-chain amino acid transport system | Non-synonymous | 411 | Ala | Gly |
| BN118_0965 | putative permease component of<br>branched-chain amino acid transport system | Non-synonymous | 574 | Met | Val |
| BN118_0965 | putative permease component of<br>branched-chain amino acid transport system | Synonymous | 75 | Leu | Leu |
| BN118_0964 | putative ATP-binding component of<br>branched-chain amino acid ABC transporter | Non-synonymous | 257 | Val | Ala |
| BN118_0964 | putative ATP-binding component of<br>branched-chain amino acid ABC transporter | Synonymous | 114 | Gly | Gly |
| BN118_0963 | putative ATP-binding component of ABC transporter | Non-synonymous | 152 | Arg | Cys |
| BN118_0961 | probable oxidoreductase | Non-synonymous | 142 | Arg | His |
| BN118_0960 | putative monooxygenase | Non-synonymous | 41 | Arg | Leu |
| BN118_0659 | conserved integral membrane protein | Synonymous | 500 | Gly | Gly |
| BN118_0189 | conserved hypothetical protein | Non-synonymous | 152 | Arg | Gly |

None of the significant Minnesota SNPs overlapped with the significant California/Washington SNPs or the historical SNPs. However, the tsr and *BN118-0965* genes were found to have SNPs in both Minnesota and California/Washington populations, though not at the same codon positions. According to the annotations in the 18323 reference genome Genbank file, BN118_0965 encodes a putative permease component of the branched-chain amino acid transport system and tsr encodes a methyl-accepting chemotaxis protein I.

### Polymorphisms characteristic of pertactin(-) strains

When all pertactin(-) strains were tested as a phenotype we identified 14 significantly associated (–log_10_P value ≥ 5) SNPs in coding regions (Table 3). Twelve of the SNPs were associated with Minnesota strains and two of the SNPs were associated with the California and Washington strains. Six of the 12 SNPs were non-synonymous.

**Table 3:** SNPs significantly associated with pertactin(-) strains. Gene and protein annotations are from the 18323 reference genome Genbank file.

| Gene | Protein | SNP Type | Codon position | Reference amino acid | Non-reference amino acid |
| --- | --- | --- | --- | --- | --- |
| BN118_0189 | conserved hypothetical protein | Non-synonymous | 152 | Arg | Gly |
| BN118_2561 | hypothetical protein, nicking endonuclease | Non-synonymous | 135 | Val | Gly |
| ctaD | cytochrome c oxidase polypeptide I | Synonymous | 420 | Lys | Lys |
| maeB | NADP-dependent malic enzyme | Non-synonymous | 628 | Gly | Asp |
| maeB | NADP-dependent malic enzyme | Non-synonymous | 628 | Gly | Ser |
| maeB | NADP-dependent malic enzyme | Non-synonymous | 625 | Gln | Arg |
| maeB | NADP-dependent malic enzyme | Synonymous | 640 | Asn | Asn |
| tsr | methyl-accepting chemotaxis protein I | Non-synonymous | 498 | Ala | Val |
| tsr | methyl-accepting chemotaxis protein I | Synonymous | 510 | Ala | Ala |
| tsr | methyl-accepting chemotaxis protein I | Synonymous | 509 | Ala | Ala |
| wbpO | capsular polysaccharide biosynthesis protein | Synonymous | 19 | Gly | Gly |
| wbpO | capsular polysaccharide biosynthesis protein | Synonymous | 20 | Tyr | Tyr |
| wbpO | capsular polysaccharide biosynthesis protein | Synonymous | 18 | Leu | Leu |
| wbpO | capsular polysaccharide biosynthesis protein | Synonymous | 21 | Val | Val |

### Polymorphisms characteristic of the pre-, whole cell and acellular vaccine periods

When all strains isolated during the acellular vaccine period (1998-2012, 62 isolates) were tested as a phenotype we identified a single significantly associated (–log_10_P value ≥ 5) non-synonymous SNP in the *BN118-0189* gene, which encodes for a restriction endonuclease. This SNP is also associated with the pertactin(-) phenotype, specifically the California/Washington pertactin(-) strains. No significant SNPs were identified from the pre-vaccine (< 1946, 3 isolates) and whole cell vaccine (1946-1997, 10 isolates) phenotypes.

## Discussion

*Bordetella pertussis* produces multiple toxins in addition to other virulence factors that facilitate within-host survival by manipulating many aspects of the human immune system [14]. The majority of comparative genomics studies of *B. pertussis* focus exclusively on these known toxins and virulence factors. Our study was motivated by the importance of understanding all genes undergoing selective pressure, not just known virulence genes. Our results did detect SNPs in known toxin and virulence genes, however the putative non-virulence SNPs we describe here are associated with outbreak isolates at a much higher level of statistical significance and therefore merit reporting. More than half of the SNPs reside in genes whose protein products are not fully characterized and whose roles in pathogenesis are unknown. Some are likely housekeeping genes without a role in pathogenesis, but others, like *BN118-0462* that is associated with lipopolysaccharide assembly, are almost certainly involved in pathogenesis.

Multiple studies indicate that *B. pertussis* relies on iron uptake systems for growth in vivo [15]. In *B. pertussis*, outer membrane receptors are required for transfer of iron chelates to the periplasm, followed by transport to the cytoplasm by ATP-binding cassette (ABC) transporters [15]. Bacterial ABC transporters are an important class of transmembrane transporters known to influence many cellular processes including antibiotic resistance, nutrient acquisition, adhesion, protein secretion and have been shown to be important for the virulence of a range of bacterial pathogens [16]. The GWAS of combined California/Washington isolates identified SNPs in both an iron uptake receptor gene, bfrI [17], and ABC transporter genes *BN118-2154, BN118-0963, BN118-0964* and *BN118-0965*. The Minnesota GWAS also identified a gene with putative iron transport functionality, *BN118-0794*, as well as multiple ABC transporter genes: *BN118-0138, BN118-2781, BN118-2428*, and *BN118.0965. BN118-0964* and *BN118-0965* are both involved in branch chain amino acid transport and *BN118-0965* had SNPs in both California/Washington and Minnesota populations, though not at the same codon positions. Branch chain amino acid transporters have been found essential for disease pathogenesis in some bacteria [18].

Another gene that contained SNPs detected in both Minnesota and California/Washington populations, but at different codon positions, was the tsr gene, which encodes a chemotaxis protein that recognizes the amino acid serine [19]. A total of six SNPs in the Minnesota strains were detected in the maeB gene that encodes NADP-dependent malic enzyme, a protein involved in pyruvate metabolism and carbon fixation [20]. Multiple SNPs in the maeB gene have been previously reported and postulated to be important in *B. pertussis* adaptation [21].

In summary, GWAS of the outbreak phenotypes identified several SNPs in genes that encode iron uptake, transporter, metabolism and chemotaxis proteins associated with in-host survivability and virulence that overlapped outbreaks, and other genes that were specific to a particular outbreak. Compared to 18323, the elevated SNPs/genome of the Minnesota outbreak strains (2408 SNPs/genome) and the nearly identical CA_2010_3‐ (7 SNPs), CA_2010_5‐ (22 SNPs) and CA_2010_4 (22 SNPs) strains suggest that subpopulations of *B. pertussis* are undergoing differential selective pressure.

Pertactin(-) *B. pertussis* strains have been increasingly reported in the U.S. and several different genotypes have been attributed to the pertactin(-) phenotype [13]. The pertactin(-) phenotypes in the strains we studied also included multiple genotypes. The GWAS of the pertactin(-) phenotype identified non-synonymous SNPs in the maeB and tsr genes of the Minnesota strains and in the *BN118-2561* and *BN118-0189* genes of the California/Washington strains. The *BN118-2561* and *BN118-0189* genes encode putative restriction endonucleases. Considering the multiple genotypes for the pertactin(-) phenotype, and the dissimilarity between maeB, tsr, *BN118.2561* and *BN118-0189* genes, these are likely confounding SNPs and not correlated or compensatory to the pertactin(-) phenotype in any way. While these SNPs may be confounding SNPs associated with the pertactin(-) phenotype, it is possible that since all the pertactin(-) strains were isolated during the acellular vaccine period that the SNPs are involved with acellular vaccine evasion and adaptation. In fact, the SNP in the *BN118-0189* gene was found to be significantly associated with the acellular vaccine period (1998-2012). No significant SNPs were identified from the pre-vaccine (<1946, 3 isolates) and whole cell vaccine (1946-1997, 10 isolates) phenotypes. While this may be due to not having enough samples for each phenotype, it is possible that the whole cell vaccine successfully blocked *B. pertussis* transmission to such an extent that evolutionary opportunities were not available for the pathogen to adapt and establish consensus strains able to evade the vaccine [22]. Analyzing additional samples should strengthen these results.

The Minnesota outbreak is alarming for several reasons. The strains isolated had higher SNPs/genome counts than all other strains, all isolates were pertactin(-), and *B. parapertussis* was also isolated (although not sequenced) [10]. The same region in Minnesota also experienced a *B. parapertussis* outbreak in 2014 [23]. This may signal the beginning of a “new normal” for *B. pertussis* and *B. parapertussis* resurgence. There are a handful of *B. parapertussis* genome sequences publicly available but more are needed for a useful GWAS and comparison to *B. pertussis*.

In a time when whole genome sequencing is becoming rapidly cheaper, the current *B. pertussis* surveillance, diagnostic and molecular typing techniques are found severely lacking. The CDC does not publish any PCR assay protocols or standards for pertussis detection, multilocus sequence typing (MLST) and multilocus variable number tandem repeat analysis (MLVA) assays only address a tiny fraction of the genome and pulsed-field gel electrophoresis (PFGE) has been found to be more discerning than MLST and MLVA combined [9]. In fact, PFGE typing suggests that pertussis isolates are evolving more rapidly on a genomic scale than in the few genes and repeat regions targeted by MLVA and MLST [9]. The GWAS we performed is precisely one way to uncover the strain diversity residing in the rest of the pertussis genome missed by MLST and MLVA assays and make more definitive conclusions about correlation. As such, it has become obvious to us that whole genome sequencing technology provides the means with which to replace the lack of a unified pertussis typing scheme and an opportunity to create a universal format for recording and reporting the molecular epidemiology of *B. pertussis* isolates that allows for more efficient comparisons between epidemics and countries.

To our knowledge, the findings reported here comprise the first GWAS, SNP count analysis and phylogenetic tree construction using whole genome *B. pertussis* sequences from strains isolated in the U.S. We are confident that additional GWAS will produce significant insights into *B. pertussis* resurgence and play an important role in surveillance diagnostics and vaccine design when applied to whole genome sequences of clinical samples in the future.

## Acknowledgements

BMA and SVS acknowledge support from the Omidyar Group and the Santa Fe Institute.

## References

[1] J. D. Cherry, “Epidemic Pertussis in 2012 The Resurgence of a Vaccine-Preventable Disease,” New England Journal of Medicine, vol. 367, no. 9, pp. 785–787, Aug. 2012. [Online]. Available: http://www.nejm.org/doi/abs/10.1056/NEJMp1209051

[2] “Pertussis | Surveillance Trend Reporting and Case Definition | CDC.” [Online]. Available: http://www.cdc.gov/pertussis/surv-reporting.html

[3] “Pertussis Epidemic California, 2014.” [Online]. Available: http://www.cdc.gov/mmwr/preview/mmwrhtml/mm6348a2.htm

[4] A. van der Zee, C. Agterberg, M. Peeters, J. Schellekens, and F. Mooi, “Polymerase Chain Reaction Assay for Pertussis: Simultaneous Detection and Discrimination of Bordetella pertussis and Bordetella parapertussis,” JOUNAL OF CLINICAL MICROBIOLOGY, vol. 31, no. 8, pp. 2134–2140, Aug. 1993.

[5] “Pertussis | Whooping Cough | Use of PCR for Diagnosis | CDC.” [Online]. Available: http://www.cdc.gov/pertussis/clinical/diagnostic-testing/diagnosis-pcr-bestpractices.html

[6] I. H. M. van Loo, K. J. Heuvelman, A. J. King, and F. R. Mooi, “Multilocus Sequence Typing of Bordetella pertussis Based on Surface Protein Genes,” Journal of Clinical Microbiology, vol. 40, no. 6, pp. 1994–2001, Jun. 2002. [Online]. Available: http://jcm.asm.org/cgi/doi/10.1128/JCM.40.6.1994-2001.2002

[7] L. M. Schouls, H. G. J. van der Heide, L. Vauterin, P. Vauterin, and F. R. Mooi, “Multiple-Locus Variable-Number Tandem Repeat Analysis of Dutch Bordetella pertussis Strains Reveals Rapid Genetic Changes with Clonal Expansion during the Late 1990s,” Journal of Bacteriology, vol. 186, no. 16, pp. 5496–5505, Aug. 2004. [Online]. Available: http://jb.asm.org/cgi/doi/10.1128/JB.186.16.5496-5505.2004

[8] K. Winter, K. Harriman, J. Zipprich, R. Schechter, J. Talarico, J. Watt, and G. Chavez, “California Pertussis Epidemic, 2010,” The Journal of Pediatrics, vol. 161, no. 6, pp. 1091–1096, Dec. 2012. [Online]. Available: http://linkinghub.elsevier.com/retrieve/pii/S0022347612005586

[9] K. E. Bowden, M. M. Williams, P. K. Cassiday, A. Milton, L. Pawloski, M. Harrison, S. W. Martin, S. Meyer, X. Qin, C. DeBolt, A. Tasslimi, N. Syed, R. Sorrell, M. Tran, B. Hiatt, and M. L. Tondella, “Molecular Epidemiology of the Pertussis Epidemic in Washington State in 2012,” Journal of Clinical Microbiology, vol. 52, no. 10, pp. 3549–3557, Oct. 2014. [Online]. Available: http://jcm.asm.org/cgi/doi/10.1128/JCM.01189-14

[10] A. G. Theofiles, S. A. Cunningham, N. Chia, P. R. Jeraldo, D. J. Quest, J. N. Mandrekar, and R. Patel, “Pertussis Outbreak, Southeastern Minnesota, 2012,” Mayo Clinic Proceedings, vol. 89, no. 10, pp. 1378–1388, Oct. 2014. [Online]. Available: http://linkinghub.elsevier.com/retrieve/pii/S0025619614007204

[11] T. J. Treangen, B. D. Ondov, S. Koren, and A. M. Phillippy, “The Harvest suite for rapid core-genome alignment and visualization of thousands of intraspecific microbial genomes,” Genome Biol, vol. 15, no. 524, p. 2, 2014. [Online]. Available: http://www.biomedcentral.com/content/pdf/s13059-014-0524-x. pdf

[12] S. G. Earle, C.-H. Wu, J. Charlesworth, N. Stoesser, N. C. Gordon, T. M. Walker, D. A. Clifton, E. G. Smith, N. Ismail, M. J. Llewelyn, and others, “Controlling for population structure in bacterial association studies,” arXiv preprint arXiv:1510.06863, 2015. [Online]. Available: http://arxiv.org/abs/1510.06863

[13] L. C. Pawloski, A. M. Queenan, P. K. Cassiday, A. S. Lynch, M. J. Harrison, W. Shang, M. M. Williams, K. E. Bowden, B. Burgos-Rivera, X. Qin, N. Messonnier, and M. L. Tondella, “Prevalence and Molecular Characterization of Pertactin-Deficient Bordetella pertussis in the United States,” Clinical and Vaccine Immunology, vol. 21, no. 2, pp. 119–125, Feb. 2014. [Online]. Available: http://cvi.asm.org/cgi/doi/10.1128/CVI.00717-13

[14] D. d. Gouw, P. W. M. Hermans, H. J. Bootsma, A. Zomer, K. Heuvelman, D. A. Diavatopoulos, and F. R. Mooi, “Differentially Expressed Genes in Bordetella pertussis Strains Belonging to a Lineage Which Recently Spread Globally,” PLoS ONE, vol. 9, no. 1, p. e84523, Jan. 2014. [Online]. Available: http://dx.plos.org/10.1371/journal.pone.0084523

[15] T. J. Brickman, C. A. Cummings, S.-Y. Liew, D. A. Relman, and S. K. Armstrong, “Transcriptional Profiling of the Iron Starvation Response in Bordetella pertussis Provides New Insights into Siderophore Utilization and Virulence Gene Expression,” Journal of Bacteriology, vol. 193, no. 18, pp. 4798–4812, Sep. 2011. [Online]. Available: http://jb.asm.org/cgi/doi/10.1128/JB.05136-11

[16] H. S. Garmory and R. W. Titball, “ATP-Binding Cassette Transporters Are Targets for the Development of Antibacterial Vaccines and Therapies,” Infection and Immunity, vol. 72, no. 12, pp. 6757–6763, Dec. 2004. [Online]. Available: http://iai.asm.org/cgi/doi/10.1128/IAI.72.12.6757-6763.2004

[17] P. Cornelis and S. C. Andrews, Eds., Iron uptake and homeostasis in microorganisms. Norfolk, UK: Caister Academic Press, 2010.

[18] S. Basavanna, S. Khandavilli, J. Yuste, J. M. Cohen, A. H. F. Hosie, A. J. Webb, G. H. Thomas, and J. S. Brown, “Screening of Streptococcus pneumoniae ABC Transporter Mutants Demonstrates that LivJHMGF, a Branched-Chain Amino Acid ABC Transporter, Is Necessary for Disease Pathogenesis,” Infection and Immunity, vol. 77, no. 8, pp. 3412–3423, Aug. 2009. [Online]. Available: http://iai.asm.org/cgi/doi/10.1128/IAI.01543-08

[19] K. K. Gosink, Y. Zhao, and J. S. Parkinson, “Mutational Analysis of N381, a Key Trimer Contact Residue in Tsr, the Escherichia coli Serine Chemoreceptor,” Journal of Bacteriology, vol. 193, no. 23, pp. 6452–6460, Dec. 2011. [Online]. Available: http://jb.asm.org/cgi/doi/10.1128/JB.05887-11

[20] I. Ganesh, S. Ravikumar, S. J. Park, S. H. Lee, and S. H. Hong, “Expression characteristics of the maeA and maeB genes by extracellular malate and pyruvate in Escherichia coli,” Korean Journal of Chemical Engineering, vol. 30, no. 7, pp. 1443–1447, Jul. 2013. [Online]. Available: http://link.springer.com/10.1007/s11814-013-0061-4

[21] M. J. Bart, S. R. Harris, A. Advani, Y. Arakawa, D. Bottero, V. Bouchez, P. K. Cassiday, C.-S. Chiang, T. Dalby, N. K. Fry, M. E. Gaillard, M. van Gent, N. Guiso, H. O. Hallander, E. T. Harvill, Q. He, H. G. J. van der Heide, K. Heuvelman, D. F. Hozbor, K. Kamachi, G. I. Karataev, R. Lan, A. Luty ska, R. P. Maharjan, J. Mertsola, T. Miyamura, S. Octavia, A. Preston, M. A. Quail, V. Sintchenko, P. Stefanelli, M. L. Tondella, R. S. W. Tsang, Y. Xu, S.-M. Yao, S. Zhang, J. Parkhill, and F. R. Mooi, “Global Population Structure and Evolution of Bordetella pertussis and Their Relationship with Vaccination,” mBio, vol. 5, no. 2, pp. e01074-14–e01074-14, Apr. 2014. [Online]. Available: http://mbio.asm.org/cgi/doi/10.1128/mBio.01074-14

[22] B. M. Althouse and S. V. Scarpino, “Asymptomatic transmission and the resurgence of bordetella pertussis,” BMC medicine, vol. 13, no. 1, p. 146, 2015.

[23] “Bordetella parapertussis Outbreak in Southeastern Minnesota in 2014.” [Online]. Available: http://www.asm.org/index.php/asm-newsroom2/press-releases/93691-bordetella-parapertussis-outbreak-in-southeastern-minnesota-in-2014

